# Use of optimized single-cell RNA flow cytometry protocol identifies monocytes as main producers of type I interferon in mouse syngeneic tumors

**DOI:** 10.1101/2024.07.23.604694

**Authors:** Khiem C. Lam, Quanyi Chen, Romina S. Goldszmid

**Affiliations:** Inflammatory Cell Dynamics Section, Laboratory of Integrative Cancer Immunology (LICI), Center for Cancer Research (CCR), National Cancer Institute (NCI), Bethesda, MD 20892, USA; Computational Biology, Bioinformatics and Genomics, Biological Sciences, University of Maryland, College Park, MD 20742, USA; Kelly Government Solutions, Bethesda, MD 20892, USA

## Abstract

The tumor microenvironment (TME) consists of complex interactions between cellular and extracellular components, among which the immune system is known to play an integral role in disease progression and response to therapy. Cytokines and chemokines are cell signaling proteins used by immune cells to communicate with each other as well as with other cell types in the body. These proteins control systemic and local immune responses and levels of cytokines/chemokines in the TME have been associated with tumor outcomes. However, cytokines and chemokines have varied expression across cell types, tumors, and host conditions. Therefore, approaches to effectively study the production of these proteins at the single-cell level in the TME are needed to fully elucidate the mechanisms governing the anti-cancer immune response. Here, we detail a protocol to assess the production of cytokines/chemokines across leukocyte populations in mouse tumors using RNA flow cytometry. Importantly, this method can be adapted with minimal changes to study various mouse and human tumors, other RNA analytes, and non-tumor tissues. With this approach, we characterize single-cell production of *Ifnb1, Xcl1* and *Ccl5* in mouse tumors and identify monocytes and monocyte-derived macrophages as the main producers of type I interferon transcript *Ifnb1* consistent across 4 different syngeneic tumor models.

## 1 Introduction

Tumors grow in an intricate microenvironment composed of cellular (e.g., tumor, immune, stroma) and extracellular (e.g., extracellular matrix, metabolites, proteins) components termed the tumor microenvironment (TME) (de Visser & Joyce, 2023). Among the cellular components, immune cells have long been implicated as key controllers of the anti-tumor response, which has led to the development of therapies targeting specific immune cells or pathways (Binnewies et al., 2018). Cytokines are cell signaling proteins largely used by the immune system to control the extent and type of immune response leading to tumor progression or resistance (Berraondo et al., 2019; Propper & Balkwill, 2022). A subset of cytokines called chemokines, or chemotactic cytokines, are important for cell migration and localization to the tumor (Berraondo et al., 2019; Propper & Balkwill, 2022). Although associated with specific tumor states, cytokine levels can vary across time, tissues, tumor types, individuals, and even gut microbiome conditions (Francescone et al., 2014; Kartikasari et al., 2021; Lam et al., 2021; Landskron et al., 2014). Moreover, the cell type(s) producing each cytokine can differ depending on these factors, leading to complex cytokine dynamics. Therefore, methodologies that can be used to readily assess the production of cytokines across cell types in the TME are needed to fully understand the anti-tumor immune responses.

Cytokine measurements are often performed using immunoassays such as enzyme-linked immunosorbent assays (ELISA) (Chiswick et al., 2012). While good for bulk measurements of homogenous samples (e.g., plasma/serum), ELISAs performed on heterogenous tissues such as the tumor lacks granularity and is confounded by factors such as variation in the proportion/number of different tumor-infiltrating cell types. Traditional flow cytometry using fluorochrome-conjugated antibodies can work well to measure certain cytokines at the single-cell level (Lovelace & Maecker, 2018; Qiu et al., 2014), but is limited to targets that are expressed at high levels and have commercially available antibodies (or custom antibodies must be generated). Single-cell RNA sequencing (scRNAseq) is a powerful tool to characterize different cell types and their state in the TME (Zhao et al., 2023). Nevertheless, using this technique to characterize intratumoral cytokines/chemokines can present several challenges: complexity of analysis, difficulties in detecting lowly expressed transcripts often leading to a reliance on cytokine signatures rather than cytokine transcripts directly, constraints in the number of cells processed which may result in potentially missed rare populations, and high costs that restrict the number of biological replicates and its use across multiple experiments and/or conditions.

RNA flow cytometry is a technique that combines flow cytometry with the in-situ hybridization approach typically used in microscopy for RNA targets, in order to quantify the expression of RNA analytes at the single-cell level (Soh & Wallace, 2018). This protocol uses the branched DNA technology employed by PrimeFlow™. In brief, permeabilized cells are hybridized with a target oligonucleotide specific to the RNA transcript of interest, followed by multiple steps of signal amplification, and hybridization with an oligonucleotide conjugated to a fluorescent dye to form branched DNA structures. The advantage of this approach is that it can be combined with standard flow cytometry practices of cell labeling with surface and intracellular fluorochrome-conjugated antibodies to not only identify cell types of interest, but also quantify their expression of RNA targets. The added benefit of this method is the specificity, high signal-to-noise ratio, relatively low costs compared to scRNAseq, and ability to assay many cells (>1 million) per sample. Multiple studies have previously used this approach to study RNA targets in the TME (Pacella et al., 2018), including cytokines/chemokines (Akeus et al., 2018; Bottcher et al., 2018; Lam et al., 2021; Ponzetta et al., 2019).

In this report, we detail a protocol (Figure 1) to quantify transcripts for the type I interferon (IFN) cytokine IFN-β and dendritic cell-attracting chemokines XCL1 and CCL5 in major immune cell types from murine tumors. We have previously employed this approach to study the anti-tumor innate immune response under different gut microbiota conditions (Lam et al., 2021). Here, we further assess the production of these molecules across tumor-infiltrating leukocyte populations. We identify monocytes and monocyte-derived macrophages as the top producers of type I IFN across syngeneic models of lymphoma, colon carcinoma, melanoma, and mammary carcinoma, which is consistent with our previous findings (Lam et al., 2021). Furthermore, we describe tumor processing techniques and considerations for tissues with high amounts of debris and dead cells. We also provide recommendations to adapt the PrimeFlow™ RNA Assay to minimize cell loss and to be able to use a conventional thermocycler when a specialized incubator is not available. Importantly, this protocol can be easily adapted to study diverse mouse and human tumors, other RNA analytes, and non-tumor tissues.

**Figure 1.**
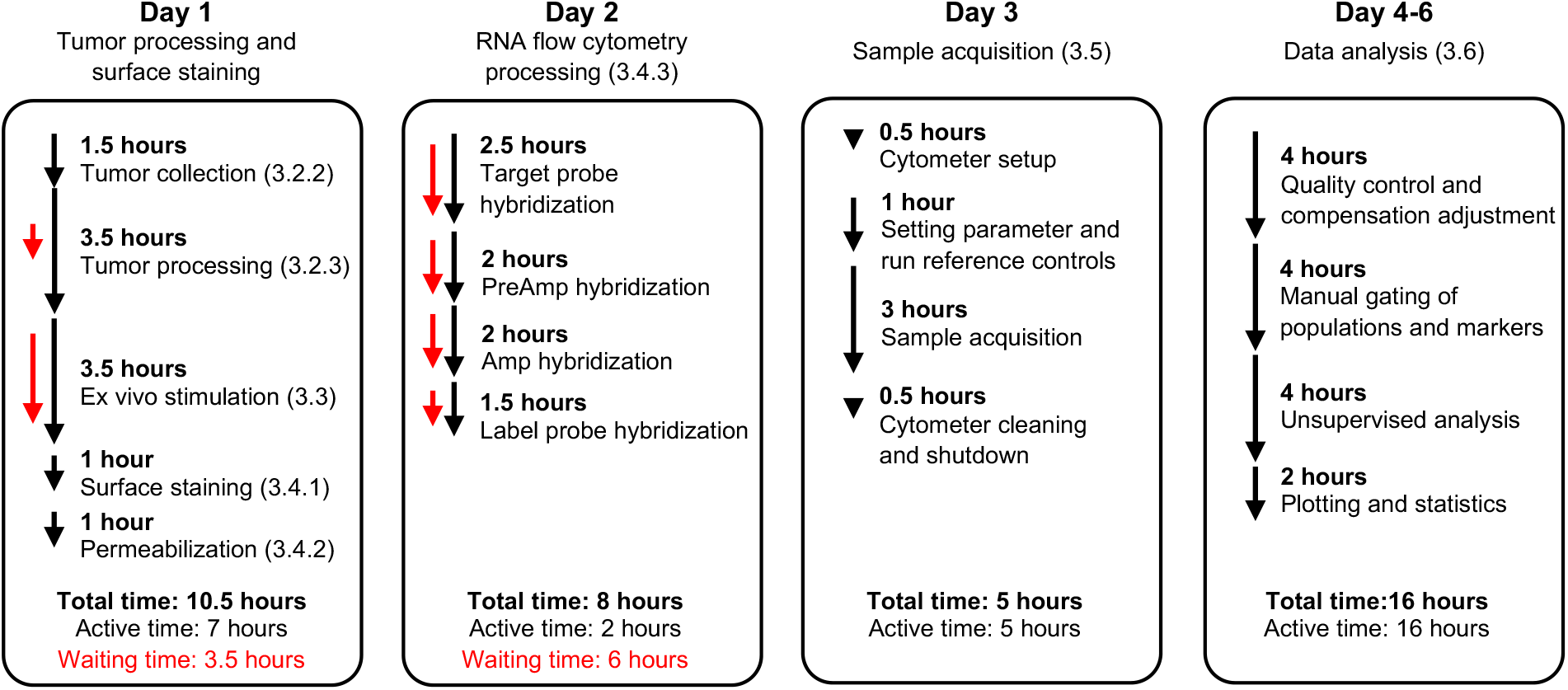
Schematic of tumor RNA flow cytometry protocol with time estimates based on experience processing 16 samples by a single person. Total times for each indicated step shown by black line while waiting/hands-off time is indicated by red line. Times will vary depending on number/complexity of samples and user expertise.

## 2 Materials

### 2.1 Common disposables

- Dissection instruments: forceps, scissors
- Serological pipettes and automatic pipettes
- Pipette tips and micropipette
- No. 20 scalpel (#371620, Bard-Parker) (see Note 1)
- 50 mL tube (#62.547.205, Sarstedt) (see Note 2)
- 60 mm petri dish (#353002, Falcon) (see Note 1)
- 40 µm cell strainer (#22-363-547, Fisher Scientific) (see Note 1)
- 3 mL syringe (#309657, BD Biosciences) (see Note 1)
- Flat-bottom 96-well plate (#167008, Thermo Scientific) (see Note 1)
- V-bottom 96-well plate (#249570, Thermo Scientific) (see Note 1)
- U-bottom 96-well plate (#163320, Thermo Scientific) (see Note 1)
- Reagent reservoir (#3054-2012, VitsaLab) (see Note 1)
- 0.2 mL 8-strip PCR tubes (#PC7061, Alkali Scientific) (see Note 1)
- Cluster tubes (#4401, Corning) (see Note 1)
- 5 mL round bottom test tubes (#352052, Corning) (see Note 1)
- 96-well 40 µm filter plate (#MANMN4010, MilliporeSigma) (see Note 1)
- 50 µM mesh (#SEFA03-50/31, Sefar) (see Note 1)
- 5 mL test tubes with cell-strainer caps (#352235, Falcon) (see Note 1)
- Enzo™ enzymatic detergent solution (#2252, Advanced Sterilization Products) (see Note 1)

### 2.2 Tissue and cell processing reagents

- RPMI 1640 medium (RPMI) (#11875093, Gibco) (see Note 1)
- DMEM medium (#10564011, Gibco) (see Note 1)
- Fetal bovine serum (FBS) (#S12450, R&D Systems) (see Note 1)
- DNase I (#10104159001, Roche) (see Note 1, Note 14)
- Collagenase IV (#17104019, Gibco) (see Note 1, Note 14)
- Phosphate buffered solution (PBS), 1X (#21-040-CV, Corning) (see Note 1)
- EDTA, 0.5 M (#E177, Amresco) (see Note 1)
- 70% ethanol (#P111000140CSGL, Thermo Fisher Scientific) (see Note 1)
- ACK lysing buffer (#118-156-101, Quality Biological)
- Trypan blue solution, 0.4% (#15250061, Gibco) (see Note 1)
- Sodium pyruvate, 100 mM (#116-079-721, Quality Biological)
- L-glutamine, 200 mM (#BEBP17-605E, Lonza) (see Note 1)
- Non-essential amino acids solution (NEAA), 100X (#11140050, Gibco) (see Note 1)
- 2-mercaptoethanol, 1000X, 55 mM (#21985023, Gibco) (see Note 1)
- Bis-(3’-5’)-cyclic dimeric adenosine monophosphate (c-di-AMP) (#tlrl-nacda, InvivoGen) (see Note 3, 22)
- RNase-free water (#351-068-721, Quality Biological) (see Note 1)

### 2.3 Flow cytometry reagents

- LIVE/DEAD™ Fixable Blue Dead Cell Stain Kit (LD Blue) (#L34962, Invitrogen)
- BD Horizon™ Brilliant Stain Buffer (#566349, BD Biosciences)
- Surface or intracellular fluorochrome-conjugated antibodies for cell identification (see Note 4, Table 1).
- PrimeFlow™ RNA Assay Kit (#88-18005-204, Invitrogen)
- Type 1 probe for Xcl1 AF647 (#VB1-19578-PF, Thermo Fisher Scientific) (see Note 5)
- Type 6 probe for Ccl5 AF750 (#VB6-14424-PF, Thermo Fisher Scientific) (see Note 5)
- Type 10 probe for Ifnb1 AF568 (#VB10-3282108-PF, Thermo Fisher Scientific) (see Note 5)
- UltraComp eBeads™ Compensation Beads (#01-2222-41, Invitrogen) (see Note 6)
- ArC™ Amine Reactive Compensation Bead Kit (#A10346, Invitrogen)
- Paraformaldehyde aqueous solution (PFA), 16% (#50-980-487, Fisher Scientific) (see Note 1, 41)

**Table 1.** Example flow cytometry panel to identify main subsets of intratumoral leukocytes by surface protein expression (see Note 4).

| Marker | Cell Target | Fluorochrome | Dilution (1/) | Manufacturer | Catalog # |
| --- | --- | --- | --- | --- | --- |
| CD11b | Myeloid cells | BUV805 | 400 | BD Biosciences | 741934 |
| CD45.2 | Leukocytes | BUV737 | 100 | BD Biosciences | 612778 |
| CD24 | Dendritic cells | BUV563 | 200 | BD Biosciences | 749336 |
| Ly6G | Neutrophils | BUV396 | 100 | BD Biosciences | 563978 |
| CD64 | Monocytes/<br>macrophages | BV786 | 100 | BD Biosciences | 741024 |
| SiglecF | Eosinophils | BV750 | 200 | BD Biosciences | 740557 |
| F4/80 | Macrophages | BV711 | 300 | BioLegend | 123147 |
| NK1.1 | NK cells | BV650 | 100 | BioLegend | 108736 |
| Ly6C | Monocytes | BV605 | 100 | BioLegend | 128036 |
| MHCII (IA/IE) | Antigen presenting cells | BV480 | 400 | BD Biosciences | 566086 |
| CD103 | Type 1 conventional dendritic cells | BV421 | 100 | BD Biosciences | 562771 |
| CD19 | B cells | PerCP-Cy5.5 | 200 | BioLegend | 115534 |
| SiglecH | Plasmacytoid dendritic cells | FITC | 100 | BioLegend | 129603 |
| CD11c | Dendritic cells | PE-Cy5.5 | 300 | Thermo Fisher Scientific | 35-0114-82 |
| CD3e | T cells | PE-Cy5 | 200 | Thermo Fisher Scientific | 15-0031-83 |
| TCRβ | α/β T cells | PE-Cy5 | 200 | Thermo Fisher Scientific | 15-5961-83 |
| TCRγ/δ | γ/δ T cells | PE-Cy5 | 200 | Thermo Fisher Scientific | 15-5711-82 |
| CD16/32 |  |  | 200 | BioXCell | BE0307 |

### 2.4 Equipment

- Refrigerated benchtop centrifuge with swinging buckets and adapters for 50 mL tube and plates (Sorvall X4 Pro, Thermofisher Scientific) (see Note 1)
- Analytical balance (#QUINTIX224-1S, Sartorius) (see Note 1)
- Water bath (#WBE05A11B, PolyScience) (see Note 1)
- Orbital mixer with 30 mm tube rack (Intelli-Mixer™ RM-2L, ELMI) (see Note 7)
- Humidified cell culture chamber (e.g., Sanyo MCO-18AIC)
- Hemocytometer (#0650030, Paul Marienfeld) (see Note 1)
- Multichannel pipette (#4661070N, Thermo Fisher Scientific) (see Note 1)
- 96-well absorbance plate reader (e.g., TECAN Spark)
- Thermocycler (e.g., Applied Biosystems MiniAmp Thermal Cycler)
- Flow cytometer (e.g., BD FACSymphony, Cytek Aurora)

### 2.5 Software

- FlowJo (FlowJo, LLC)
- R and package to perform unsupervised dimensional reduction and clustering such as Cytofkit (Chen et al., 2016) or Spectre (Ashhurst et al., 2022)

## 3 Methods

### 3.1 Reagent preparation

These reagents can be prepared the day before collection/processing:

- FACS buffer (12 mL/sample; stored 4°C): PBS supplemented with 2% FBS and 2 mM EDTA
- PBS +EDTA (1 mL/sample; stored 4°C): PBS supplemented with 2 mM EDTA
- 50 mL tubes with 5 mL of RPMI supplemented with 0.5% FBS (1 tube/sample; stored 4°C)
- RPMI complete (1 mL/sample; stored 4°C): RPMI supplemented with 10% FBS, 1 mM sodium pyruvate, 55 µM 2-mercaptoethanol, 0.1 mM NEAA, and 2 mM L-glutamine

### 3.2 Tumor collection and processing

#### 3.2.1 Generation of in vivo tumors

1. Generate in vivo mouse tumors by implanting mouse cancer cell lines in the desired anatomical location (e.g., 5 × 10^5^ EL4 Lymphoma cells implanted subcutaneously in the flank) (see Note 8).
2. Monitor tumor growth using caliper measurements at least 2-3 times per week or according to institutional ACUC guidelines. Calculate tumor volume (mm^3^) using:

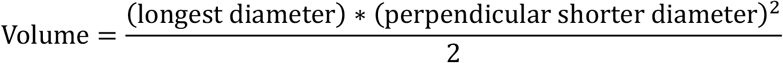
3. Collect tumors when volume is approximately 200-500 mm^3^ (see Note 9).

#### 3.2.2 Tumor collection

1. Euthanize tumor-bearing mice in accordance with institutional ACUC guidelines (see Note 10).
2. Using forceps and scissors, gently open the skin away from the tumor.
3. While folding the skin back to reveal the internal side of the tumor, identify and remove the tumor draining lymph node (e.g., inguinal lymph node for subcutaneous tumors implanted in the flank) using forceps and a scalpel if necessary (see Note 11).
4. Remove excess fat surrounding the outside of the tumor.
5. Using a scalpel, gently score around the margins of the tumor and slice under the tumor to separate it from the skin (see Note 12).
6. Measure and record tumor weight using analytical balance.
7. Place each tumor (weighing 50-200 mg) into separate 50 mL tubes containing 5 mL of RPMI supplemented with 0.5% FBS and place on ice or at 4°C (see Note 13 if tumors are >220 mg or <50 mg).

#### 3.2.3 Tumor processing

1. Pour tumor sample with media into a petri dish placed on ice. Mechanically disrupt tumors with scalpel or scissors into approximately 1-2 mm pieces.
2. Pour sample with media back into its 50 mL tube using forceps to ensure all pieces are back into the tube.
3. Using a micropipette, collect 1 mL of media from the sample tube and wash the petri dish to collect any remaining content before placing back into 50 mL tube.
4. Repeat steps 1-3 for all samples using a new petri dish and cleaning instruments with 70% ethanol between each sample to prevent cross-contamination.
5. Add 50 µL of DNase I stock (10 mg/mL) and 50 µL of Collagenase IV stock (20,000 U/mL) into each 50 mL tube (see Note 14).
6. Vortex or shake each tube for a few seconds to mix.
7. Place tubes in a water bath at 37°C for 10 minutes.
8. Transfer tubes to orbital mixer placed inside of 37°C incubator (ensure tube lids are on correctly and secured to avoid loss of sample while rotating) (see Note 15). Set mixer to rotate 135° in each direction followed by a brief shaking (i.e., Mode F8, 25 RPM). Incubate sample for 50 minutes (see Note 16).
9. Add 10 mL cold FACS buffer to each tube to stop digestion and place on ice (see Note 17).
10. Pour contents of sample tube through a 40 µm cell strainer set in a new 50 mL tube, mashing the rest of tissue that may still be intact with the back of a 3 mL syringe plunger. Pour the cells and media through the cell strainer back into the old 50 mL tube before passing through the cell strainer once again into the new tube.
11. Centrifuge sample tubes at 600 x *g* for 5 min at 4°C.
12. Aspirate and discard supernatant carefully to not disturb cell pellet.
13. Resuspend cell pellet with 1 mL of ACK lysing buffer using a micropipette. Place sample on ice to incubate for 3-5 minutes.
14. Add 10 mL of FACS buffer to each tube.
15. Centrifuge sample tubes at 500 x *g* for 5 min at 4°C.
16. Aspirate and discard supernatant carefully to not disturb cell pellet.
17. Resuspend cells in 200 µL FACS buffer and transfer into a flat-bottom 96-well plate.
18. Measure OD550 using plate reader to estimate cell density (see Note 18).
19. If absolute quantification of cell number is required, count live cell concentration using preferred technique (e.g., dead cell exclusion using Trypan blue and counting with a hemocytometer).

### 3.3 Ex vivo stimulation for detection of cytokines/chemokines

1. Place 8 × 10^6^ total cells (i.e., live, dead, cellular debris) per sample into 2 wells (for unstimulated and stimulated) of a V-bottom 96-well plate (see Note 19). Add additional wells for controls using combined leftover of tumor single-cell suspension: cytometer setup, unstained, FMO *Ifnb1*, FMO *Xcl1*, FMO *Ccl5* (see Note 20).
2. Centrifuge sample plate at 500 x *g* for 5 min at 4°C.
3. After checking for visible pellet (see Note 21), discard supernatant either by swiftly flicking plate upside down into a receptacle or aspirating manually using a multichannel pipette carefully to not disturb the cell pellet.
4. Resuspend respective samples in 200 µL RPMI complete with or without 100 µg/mL c-di-AMP (see Note 22) and transfer to U-bottom plate (see Note 23).
5. Incubate sample plate in a humidified incubator (see Note 24) at 37°C 5% CO2 for 3 hours (see Note 25).
  a. While samples are incubating, prepare surface antibody mix (50 µL per sample/well, see Note 26) in FACS buffer or Brilliant Stain Buffer depending on the fluorochromes used (see Note 4). Store surface antibody mix at 4°C in the dark until sample staining Step 3.4.1.7.
6. After 3-hour incubation, transfer samples back to a V-bottom plate and centrifuge at 500 xg for 5 min at 4°C.
7. After checking for visible pellet (see Note 21), discard supernatant either by swiftly flicking plate upside down into a receptacle or aspirating manually using a multichannel pipette carefully to not disturb the cell pellet. If desired, the supernatant can be removed and stored separately, instead of discarding, to measure any potential extracellular analytes of interest.
8. Resuspend samples in 200 µL PBS +EDTA.
9. Centrifuge sample plate at 500 x *g* for 5 min at 4°C.
10. After checking for visible pellet (see Note 21), discard supernatant either by swiftly flicking plate upside down into a receptacle or aspirating manually using a multichannel pipette carefully to not disturb the cell pellet.
11. Repeat steps 8-10 to perform an additional wash of the samples (see Note 27).

### 3.4 Flow cytometry staining

#### 3.4.1 Viability and surface staining

1. Place RNA wash buffer at room temperature to warm up (see Note 28).
2. Resuspend all samples but unstained control in 100 µL of LD Blue (see Note 30) diluted 1:1000 in PBS +EDTA (prepared in bulk 100 µL/sample, see Note 26). For unstained control, resuspend in 100 µL PBS +EDTA.
3. Incubate sample plate for 15 min at 4°C in the dark.
4. Add 100 µL of cold FACS buffer to each well.
5. Centrifuge sample plate at 500 x *g* for 5 min at 4°C.
6. After checking for visible pellet (see Note 21), discard supernatant either by swiftly flicking plate upside down into a receptacle or aspirating manually using a multichannel pipette carefully to not disturb the cell pellet.
7. Resuspend all samples but unstained control with 50 µL surface antibody mix (prepared in bulk 50 µL/sample in Step 3.3.5). For unstained control, resuspend in 50 µL of same buffer used to make antibody mix (e.g., Brilliant Stain Buffer or FACS buffer).
8. Incubate sample plate for 10 min at 4°C followed by 15 min at room temperature in the dark.
9. Add 150 µL of cold FACS buffer to each well.
10. Centrifuge sample plate at 500 x *g* for 5 min at 4°C.
  a. During centrifugation step, prepare Fixation Buffer 1 (200 µL/sample, see Note 26) by mixing equal parts PrimeFlow RNA Fixation Buffer 1A and 1B. Avoid vortexing or vigorously shaking this buffer, inverting the tube several times to mix should suffice.
11. After checking for visible pellet (see Note 21), discard supernatant either by swiftly flicking plate upside down into a receptacle or aspirating manually using a multichannel pipette carefully to not disturb the cell pellet.

#### 3.4.2 Permeabilization and intracellular staining

1. Resuspend all samples with 200 µL Fixation Buffer 1.
2. Incubate sample plate for 30 min at 4°C in the dark.
  a. While samples are incubating, prepare 1X PrimeFlow RNA Permeabilization Buffer with RNase Inhibitors by diluting PrimeFlow RNA Permeabilization Buffer (10X) to 1X with RNase-free water (200 μL/sample or 500 μL/sample if protein intracellular antibody staining is being performed in Step 8 below, see Note 26). Add RNase Inhibitor (1,000X) at a 1:000 dilution. Mix gently by inverting tube. Keep on ice or at 4^°^C.
3. Centrifuge sample plate at 1,000 x *g* for 4 min at 4^°^C.
4. After checking for visible pellet (see Note 21), aspirate and discard supernatant manually using a multi-channel pipette taking care not to disturb the cell pellet (see Note 31).
5. Resuspend all samples with 1X PrimeFlow RNA Permeabilization Buffer with RNase Inhibitors.
6. Centrifuge sample plate at 1,000 x *g* for 4 min at 4^°^C.
  a. While samples are centrifuging, prepare 1X PrimeFlow RNA Fixation Buffer 2 by diluting PrimeFlow RNA Fixation Buffer 2 (8X) with PrimeFlow RNA Wash Buffer (200 µL/sample, see Note 26). Mix gently by inverting tube. If performing protein intracellular staining in Step 8, prepare intracellular antibody mix here too.
7. After checking for visible pellet (see Note 21), aspirate and discard supernatant manually using a multi-channel pipette taking care not to disturb the cell pellet (see Note 31).
8. If performing protein intracellular staining using fluorochrome-conjugated antibodies, proceed with the following steps (see Note 32):
  a. Prepare the intracellular antibody mix using the PrimeFlow RNA Permeabilization Buffer with RNase Inhibitors (50 μL/sample, see Note 26).
  b. Resuspend cells in 50 μL of intracellular antibody mix (except unstained control that is resuspended in the same buffer used to prepare the antibody mix).
  c. Incubate for 30 min at 4^°^C in the dark.
  d. Add 150 μL of cold PrimeFlow RNA Permeabilization Buffer with RNase Inhibitors to each well.
  e. Centrifuge sample plate at 1,000 x *g* for 4 min at 4^°^C
  f. After checking for visible pellet (see Note 21), aspirate and discard supernatant manually using a multi-channel pipette taking care not to disturb the cell pellet (see Note 31).
9. Resuspend each sample in 200 μL of Fixation Buffer 2.
10. Incubate overnight at 4^°^C in the dark (see Note 33).

#### 3.4.3 RNA flow cytometry processing (see Note 34)

1. Prior to starting this section (30-60 minutes), perform the following steps:
  a. Set water bath and thermocycler to 40^°^C to warm-up.
  b. Place RNA wash buffer at room temperature to warm up (see Note 28).
  c. Thaw Target Probe Sets (20X) at room temperature.
  d. Pre-warm PrimeFlow RNA Target Probe Diluent at 40^°^C in water bath.
2. Centrifuge sample plate at 1,000 x *g* for 4 min at room temperature.
3. After checking for visible pellet (see Note 21), aspirate and discard supernatant manually using a multi-channel pipette taking care not to disturb the cell pellet (see Note 31).
4. Resuspend each sample in 200 μL of RNA wash buffer.
5. Centrifuge sample plate at 1,000 x *g* for 4 min at room temperature.
  a. While samples are centrifuging, dilute Target Probes (20X) 1:20 in RNA Target Probe diluent (50 µL/samples, see Note 26) (see Note 35). Mix by pipetting up and down.
6. After checking for visible pellet (see Note 21), aspirate and discard supernatant manually using a multi-channel pipette taking care not to disturb the cell pellet (see Note 31, 36).
7. Resuspend each sample in 50 µL PrimeFlow RNA Wash Buffer and transfer to 8-tube strip PCR tubes.
8. Add 50 µL of Target Probe mix to each sample and mix by pipetting up and down. Use PrimeFlow RNA Wash Buffer for Unstained controls and appropriate Target Probe mixes for FMO controls.
9. Incubate sample tubes in thermocycler at 40^°^C for 2 hours (see Note 37).
  a. While samples are incubating, prepare PrimeFlow RNA Wash Buffer with RNase Inhibitors by adding RNase Inhibitors (100X) to PrimeFlow™ RNA Wash Buffer at a 1:100 dilution (50 µL/sample, see Note 26). Mix gently by inverting. Pre-warm PrimeFlow RNA PreAmp Mix at 40^°^C in water bath.
10. Add 100 µL PrimeFlow RNA Wash Buffer to each tube. Mix gently by inverting tube.
11. Centrifuge sample tubes at 1,000 x *g* for 4 min at room temperature (see Note 38).
12. After checking for visible pellet (see Note 21), aspirate and discard supernatant manually using a multi-channel pipette taking care not to disturb the cell pellet (see Note 31).
13. Resuspend each sample in 200 µL PrimeFlow RNA Wash Buffer.
14. Centrifuge sample tubes at 1,000 x *g* for 4 min at room temperature (see Note 38).
15. After checking for visible pellet (see Note 21), aspirate and discard supernatant manually using a multi-channel pipette taking care not to disturb the cell pellet (see Note 31, 36).
16. Resuspend each sample in 50 µL PrimeFlow RNA Wash Buffer with RNase Inhibitors (see Note 39).
17. Add 50 µL of PrimeFlow RNA PreAmp Mix to each sample and mix by pipetting up and down.
18. Incubate sample tubes in thermocycler at 40^°^C for 1.5 hours (see Note 37).
  a. While samples are incubating, pre-warm PrimeFlow RNA Amp Mix at 40^°^C in water bath.
19. Repeat steps 10-15 to wash the samples.
20. Add 50 µL of PrimeFlow RNA Wash Buffer too each sample. Add 50 µL of PrimeFlow RNA Amp Mix to each sample and pipet to mix.
21. Incubate sample tubes in thermocycler at 40^°^C for 1.5 hours (see Note 37).
  a. While samples are incubating, pre-warm PrimeFlow RNA Label Probe Diluent at 40^°^C in water bath and place PrimeFlow RNA Label Probes (100X) at 4°C in the dark to thaw.
22. Repeat steps 10-15 to wash the samples.
  a. During centrifugation step, dilute PrimeFlow™ RNA Label Probes (100X) 1:100 in PrimeFlow RNA Label Probe Diluent (50 µL/sample, see Note 26).
23. Add 50 µL of PrimeFlow RNA Wash Buffer too each sample. Add 50 µL Label Probe mix to each sample and pipet to mix.
24. Incubate sample tubes in thermocycler at 40^°^C for 1 hour (see Note 37).
25. Repeat steps 10-15 to wash the samples.
  a. During centrifugation step, prepare mix of PrimeFlow RNA Storage Buffer and eBioscience IC Fixation Buffer at 1:1 ratio (200 µL/sample, see Note 26).
26. Resuspend each sample in 200 µL of RNA Storage Buffer/IC Fixation Buffer mix.
27. Store samples at 4°C in the dark for up to 5 days before acquisition.

#### 3.4.4 Single-color reference controls (see Note 40)

1. Make 1:10 dilutions of fluorochrome-conjugated antibodies (except compensation controls for PrimeFlow fluorochromes) in FACS buffer.
2. Vortex UltraComp eBeads™ Compensation Beads well and add 15 µL to each well of 96-well plate for each antibody. Add 85 µL of FACS buffer to each well with Ultracomp beads (except compensation controls for PrimeFlow fluorochromes).
3. Add 1 µL of diluted panel antibodies or 2 µL of reference controls for PrimeFlow fluorochromes to their respective wells (see Note 29). Pipet to mix.
4. Vortex Arc reactive beads well and add 1 drop to well for LD Blue reference control.
5. Add 1 µL of LD Blue (see Note 30) to well with Arc reactive beads. Pipet to mix.
6. Incubate reference plate for 15 min at 4°C in the dark.
7. Add 100 µL of cold FACS buffer to each well.
8. Centrifuge sample plate at 600 x *g* for 5 min at 4°C.
9. After checking for visible pellet (see Note 21), discard supernatant either by swiftly flicking plate upside down into a receptacle or aspirating manually using a multichannel pipette carefully to not disturb the pellet.
10. Resuspend each sample in 150 µL of cold FACS buffer.
11. Vortex Arc negative beads well and add 1 drop to well for LD Blue reference control.
12. Centrifuge sample plate at 600 x *g* for 5 min at 4°C.
13. After checking for visible pellet (see Note 21), discard supernatant either by swiftly flicking plate upside down into a receptacle or aspirating manually using a multichannel pipette carefully to not disturb the pellet.
14. Resuspend all reference samples in 100 µL 4% PFA (see Note 41).
15. Incubate reference plate for 10 min at room temperature in the dark.
16. Add 100 µL of FACS buffer to each well.
17. Centrifuge sample plate at 600 x *g* for 5 min at 4°C.
18. After checking for visible pellet (see Note 21), discard supernatant either by swiftly flicking plate upside down into a receptacle or aspirating manually using a multichannel pipette carefully not to disturb the pellet.
19. Resuspend all reference samples in 200 µL FACS buffer.
20. Store reference samples at 4°C in the dark until sample acquisition.

### 3.5 Sample acquisition

1. Set up cytometer according to directions provided by manufacturer or institutional facilities (see Note 42).
2. Filter samples using 40 µm 96-well filter plate (see Note 43) then transfer to appropriate acquisition tube for available cytometer (e.g., 96-well plate, cluster tubes, 5 mL test tubes) (see Note 44).
3. Acquire single-color reference controls and setup compensation/unmixing.
4. Using the “cytometer setup “ control, set FSC-A and SSC-A voltages so that the Live CD45^+^ cells are within the linear range. Make preliminary gates of major cell types with diverse FSC-A and SSC-A (e.g., eosinophils, neutrophils, T cells) to ensure correct settings.
5. Acquire samples taking care to monitor fluidics and data preview (see Note 45).
6. Export data as FCS files.
7. Clean and shutdown cytometer according to directions provided by manufacturer or institutional flow cytometry core facility.

### 3.6 Data analysis

1. Import sample FCS files into FlowJo or a similar software used to analyze flow cytometry data.
2. Perform initial gates for cells as previously described (Araya & Goldszmid, 2020):
  a. Exclude inconsistencies in sample acquisition due to fluidics by gating on the parameter in the last position of laser delay settings (e.g., YG586) vs Time.
  b. Cells: exclude debris by gating on FSC-A vs SSC-A.
  c. Singlets: exclude doublets by gating on FSC-H vs FSC-A.
  d. Live: exclude dead cells by gating on SSC-A vs LD Blue to remove LD Blue^+^.
  e. Leukocytes: exclude non-immune cells by gating on SSC-A vs CD45 to remove CD45^neg^.
3. Assess data quality and adjust compensation values for pairwise markers as needed to remove artifacts from fluorescence spillover that were not corrected properly during automated compensation removal (Araya & Goldszmid, 2020; Roederer, 2002) (see Note 46).
4. Perform manual gating of cell populations of interest based on primary identification markers.
5. Using FMO controls to define negative signal, create gates to define positive signal of RNA markers for each cell type (see Note 47).
6. Optionally, gate total positive signal of RNA markers using SSC-A vs RNA and export to perform unsupervised dimensional reduction and clustering (Araya & Goldszmid, 2020) to define the proportion of each cell type that are producing that cytokine/chemokine. This can be done in R using packages such as Cytofkit (Chen et al., 2016) or Spectre (Ashhurst et al., 2022).
7. Quantify and plot the median fluorescence intensity (MFI) indicative of expression level of each cytokine per cell type (Figure 2A-B), the proportion of cytokine-producing cells (i.e., frequency of population, frequency of leukocytes) (Figure 2C-D), and/or the total number of each cell type producing each cytokine (Figure 2E).

**Figure 2.**
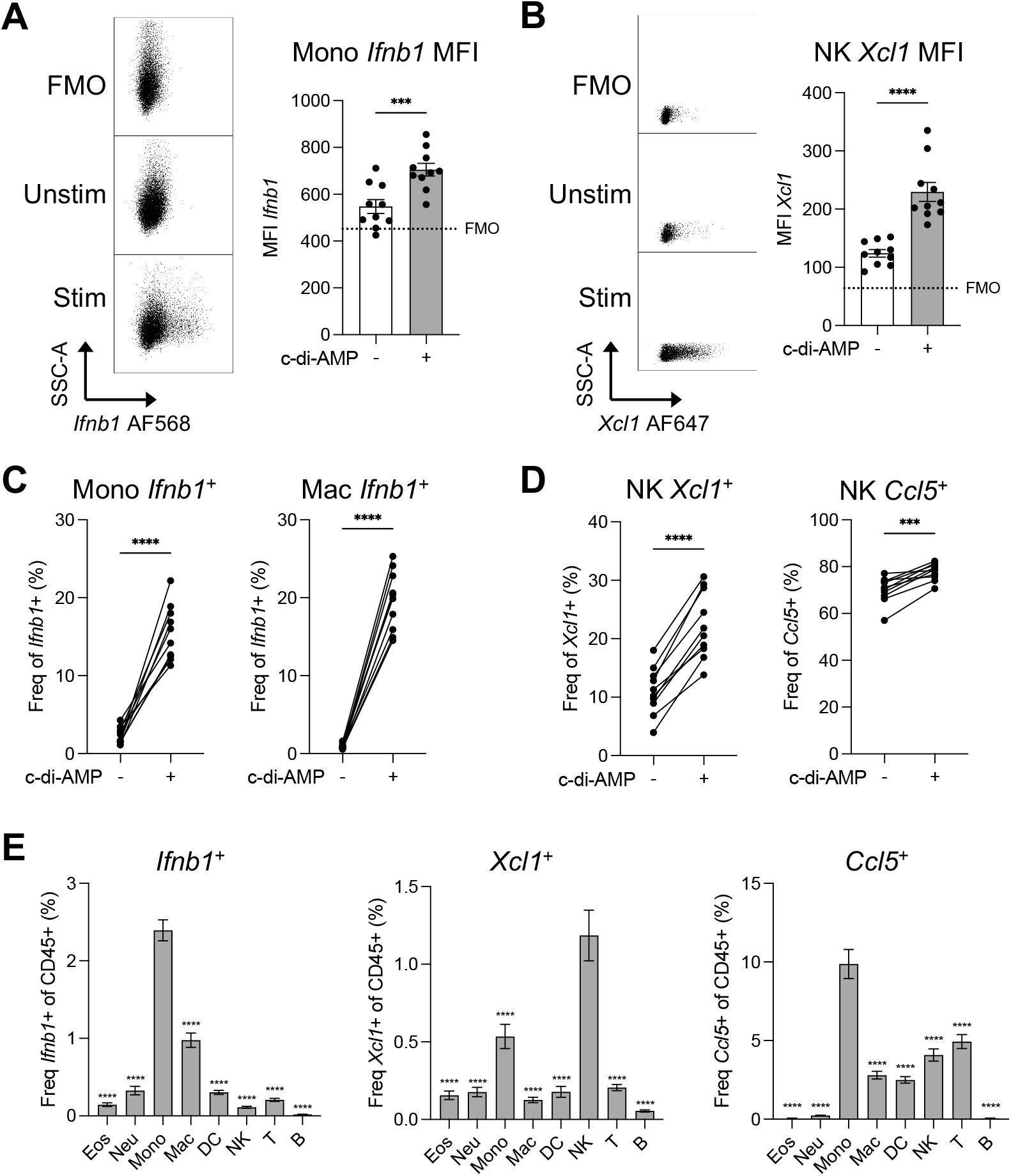
Analysis of cytokines/chemokines transcripts (*Ifnb1, Xcl1, Ccl5*) by RNA flow cytometry in EL4 lymphoma tumor-infiltrating leukocytes (Live CD45^+^) unstimulated (Unstim) or stimulated (Stim) with c-di-AMP for 3 hours ex vivo. Major leukocyte cell types are shown as eosinophils (Eos), neutrophils (Neu), monocytes (Mono), macrophages (Mac), dendritic cells (DCs), natural killer cells (NKs), T cells (T), and B cells (B). (**A**) Representative dot plot of *Ifnb1* expression in Mono across different conditions (left) and bar plot of *Ifnb1* median fluorescence intensity (MFI) in Mono (right). (**B**) Representative dot plot of *Xcl1* expression in NKs across different conditions (left) and bar plot of *Xcl1* MFI in NKs (right). (**C**) Frequency of *Ifnb1*^+^ cells of parental populations Mono (left) and Mac (right). (**D**) Frequency of *Xcl1*^+^ (left) and *Ccl5*^+^ (right) cells of total NKs. (**E**) Proportion of *Ifnb1*^+^ (left), *Xcl1*^+^ (center), and *Ccl5*^+^ cells of each major leukocyte population after c-di-AMP stimulation. Data shown as mean ± SEM with n = 10 biological replicates. Data in A-D compared using paired t-test. Data in E compared between each cell type and the highest producing cell type using one-way ANOVA with Holm-Sidak multiple hypothesis correction. ***p < 0.001, ****p < 0.0001.

## 4 Results and discussion

Here, we have developed a protocol to characterize the production of cytokines/chemokines among leukocyte populations of mouse syngeneic tumors using RNA flow cytometry. This method can be adapted with minimal variation to analyze different mouse and human tumors, other RNA targets, and non-tumor tissues (e.g., lung, liver, spleen). Moreover, this approach can be employed to address various scientific questions such as: which cell types are producing the cytokines/chemokines in the TME, how much cytokine/chemokine is produced by a particular cell type, and what are the differences in overall and cell-type specific production of cytokines/chemokines between experimental groups. We offer examples of such data in Figure 2, Figure 3, and in our previous study (Lam et al., 2021) that can be leveraged to gain mechanistic insights into the anti-tumor immune response and the identification of therapeutic targets.

**Figure 3.**
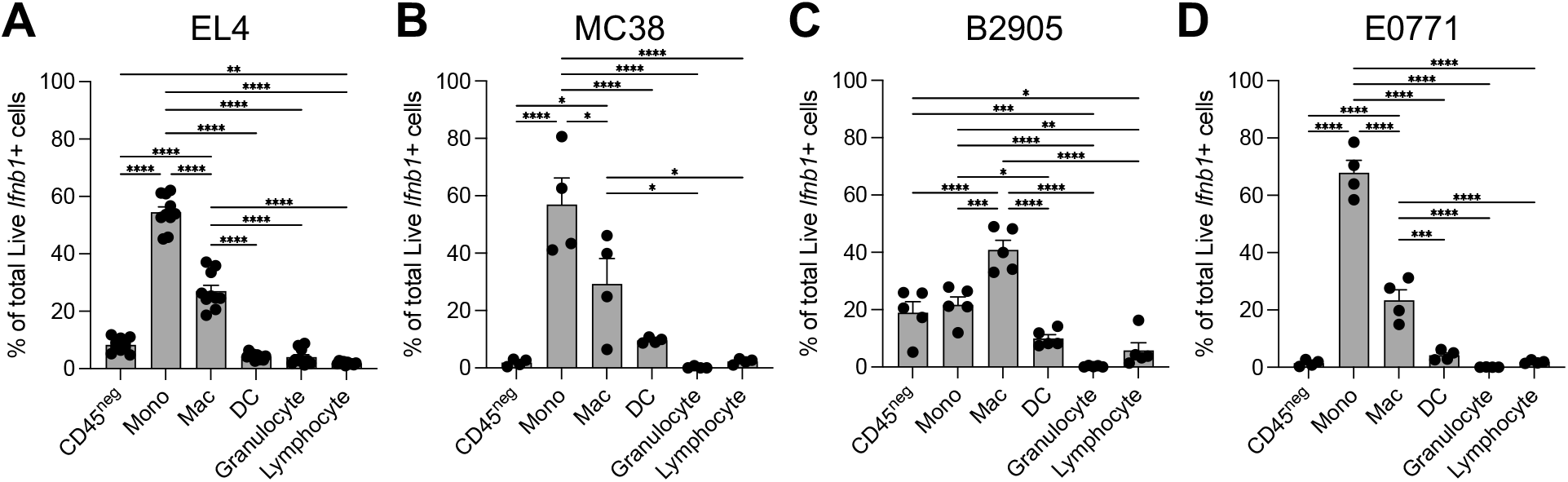
Proportion of Live *Ifnb1*^+^ tumor-infiltrating cells from C57BL/6 syngeneic mouse tumors after ex vivo stimulation with c-di-AMP for 3 hours. Major populations shown as non-immune (CD45^neg^), monocytes (Mono), macrophages (Mac), dendritic cells (DCs), granulocyte (i.e., eosinophils, neutrophils), and lymphocyte (i.e., NK, B, T cells). (**A**) EL4 lymphoma tumors collected 9 days after implantation with 5 × 10^5^ cells subcutaneously in right flank (n = 10). (**B**) MC38 colon carcinoma tumors collected 10 days after implantation with 2 × 10^5^ cells subcutaneously in right flank (n = 4). (**C**) B2905 melanoma tumors collected 34 days after implantation with 1 × 10^6^ cells subcutaneously in right flank (n = 5). (**D**) E0771 mammary tumors collected 34 days after implantation with 2 × 10^5^ cells into 4^th^ left mammary fat pad (n = 4). Comparisons between each cell type performed using one-way ANOVA with Tukey multiple hypothesis correction. *p < 0.05, **p < 0.01, ***p < 0.001, ****p < 0.0001.

Whereas monocyte *Ifnb1* expression is low in the steady-state (unstimulated) TME of EL4 lymphoma tumors (Figure 2A, C), NK cells show high expression of *Xcl1* and *Ccl5* (Figure 2B, D), and all three cytokines/chemokines increase upon c-di-AMP stimulation; this highlights the sensitivity of RNA flow cytometry and its ability to capture a variety of cytokines/chemokines and their changes across conditions. Therefore, depending on the cytokine/chemokine of interest (e.g., *Xcl1, Ccl5*), detection can be performed directly ex vivo without the need for stimulation. In response to the STING-agonist c-di-AMP, naturally produced by the gut microbiota, monocytes are the main immune producers of *Ifnb1* and *Ccl5* while NK cells are the main source of *Xcl1* (Figure 2E).

We further assessed the contribution of both immune and non-immune cells to the total production of *Ifnb1* across 4 syngeneic models (Figure 3A-D): EL4 lymphoma, MC38 colon carcinoma, B2905 melanoma, and E0771 mammary carcinoma. Remarkably, monocytes were found to be the main producers of *Ifnb1* in EL4, MC38, E0771 (Figure 3A, B, D) followed by macrophages. In B2905, macrophages were the largest producers of *Ifnb1* followed by monocytes and non-immune cells (Figure 3C). We have previously shown that the majority of macrophages in these type of syngeneic tumors are of monocytic origin (Lam et al., 2021); thus, we find monocytes and their differentiated state, monocyte-derived macrophages, are the main producers of type I IFN in the TME, consistent with our previous report (Lam et al., 2021). Although to a lesser extent than monocytes, we do observe type I IFN production by other cell types such as dendritic cells and non-immune cells in the tumor, in line with previous reports (Yu et al., 2022).

Experience with multiparametric flow cytometry, while not required, does increase accessibility and success with this technique. Furthermore, because this approach targets RNA expression, it may not be fully representative of protein expression and should be validated using other techniques such as ELISAs or cell-specific gene deletions. Despite these and potentially other limitations, this protocol offers a highly adaptable, cost-effective approach to examine the production of cytokines/chemokines across a wide array of cell types in the tumor microenvironment.

## 5 Methods and troubleshooting notes

1. Product catalog numbers and source are provided as a reference, but equivalent products from other sources can be used.
2. If using an orbital mixer to mix samples during digestion step (recommended), make sure to test 50 mL tubes beforehand to ensure there is no leakage (e.g., fill with water and place on orbital shaker for 1 hour). We have not had leakage with this specific brand of 50 mL tubes (#62.547.205, Sarstedt) but have had leakage with some other brands.
3. Depending on the cytokine/chemokine of interest, ex vivo stimulation may be required for detection. Here, we used the cyclic dinucleotide c-di-AMP, which is a known stimulator of interferon genes (STING) agonist, to induce expression of type I interferon and the chemokines XCL1 and CCL5. The molecule used for stimulation will vary depending on the context of the project, the target cells, and the cytokines/chemokines of interest. Of note, certain targets, such as *Xcl1* and *Ccl5*, can be directly detected ex vivo without stimulation.
4. Table 1 contains a list of surface markers and antibodies given as an example of a flow cytometry panel to analyze major immune cells that can be found in mouse tumors. This should be adapted to the specific needs of the project, cell types of interest, and the configuration of the available flow cytometer. See (Araya & Goldszmid, 2020) for more information regarding development and analysis of multiparametric flow cytometry panels. These surface markers are, for the most part, used in combination in order to define specific cell populations, particularly among myeloid cells.
5. Careful consideration of probe type is needed for each RNA analyte. Up to four RNA targets can be analyzed simultaneously in a single sample using different probes. As detailed by the manufacturer, Type 1 (AF647) and Type 10 (AF568) probes should be used for lowly expressed RNA targets because they are the brighter fluorochromes while Type 4 (AF488) and Type 6 (AF750) should be used for highly expressed targets because they are the dimmer fluorochromes. For example, *Ifnb1* and *Xcl1* are known to be lowly expressed transcripts so Type 1 and Type 10 probes were used, respectively, while the Type 6 probes was used for *Ccl5* which is highly expressed.
6. A bottle of the UltraComp eBeads should be provided with the PrimeFlow RNA Assay Kit. This catalog is provided in case additional amounts are needed.
7. While favorable to have an orbital mixer to aide in homogeneous digestion of tumor tissues, the protocol can be adapted if such a mixer is not available. See Note 15.
8. Other implantable and different locations (e.g., mammary fat pad; subcutaneously into the back) or genetically engineered tumor models can be used as well. In the case of implantable tumor models, the number of cells will vary depending on the model and appropriate institutional ACUC guidelines.
9. Tumors that are too small will not have enough cells to isolate and perform the analysis. However, tumors that are too large often become necrotic and ulcerated leading to many dead cells in downstream processes and increased noise in the data. Therefore, it is optimal to collect tumors in the 200-500 mm^3^ range although several smaller tumors can be pooled for processing as described in subsequent steps. The time to reach the optimal tumor volume will depend on the tumor type. For example, subcutaneous EL4 tumors would reach the optimal volume within 7-9 days post-implantation while subcutaneous MC38 colon carcinoma may take 10-12 days.
10. Animals must be euthanized by trained personnel using appropriate technique, equipment, and agents according to their institutional ACUC guidelines (e.g., CO2 inhalation).
11. It is important to remove the tumor draining lymph node as leaving them in could contaminate the tumor processing and make the results uninterpretable.
12. Be careful when performing this procedure ensuring that the scalpel is pointed away from you and is horizontal to the mouse skin to prevent injury.
13. Ideally, tumors for processing should be between 50-200 mg to have enough cells for flow cytometry, but not too much as to hinder processing. Different tumor types may have different amounts of infiltrating cells, but this is a general guideline that needs to be optimized for specific tumor models. If tumors are >220 mg, it is recommended to take a representative section (i.e., pizza slice for round tumors) of at least ¼ of the tumor to reach between 100-200 mg; weigh section using analytical balance and proceed with the protocol. If tumors are <50 mg, it is recommended to combine multiple similarly sized tumors within the same experimental group ensuring similar characteristics (e.g., mouse age, sex) to achieve between 50-200 mg and proceed with the protocol.
14. It is recommended that small aliquots of DNase I stock (10 mg/mL in PBS) and Collagenase IV stock (20,000 U/mL in DMEM) be made beforehand (e.g., 500 µL enough for 10 samples) and kept at -20°C for use. Avoid multiple freeze-thaws of these enzymes.
15. If an orbital mixer is not available, incubate samples at 37°C in a water bath vortexing for 10 seconds every 15 min during the total time of incubation.
16. The incubation time should be optimized depending on the consistency of the tumor. For EL4 and MC38 tumors which are more solid, 40-50 minutes is needed to digest the tissue. However, for tumors that are less solid (e.g., B16-F10), 30 minutes of incubation may be more appropriate.
17. It is critical to add cold buffer containing 2-10% FBS to stop enzymatic activity.
18. It is critical to normalize the amount of each sample for flow cytometry to ensure staining conditions are consistent across samples. This can be done through counting number of cells and cellular debris using a hemocytometer (Trypan Blue or other exclusionary dyes can be used to quantify proportion of live cells simultaneously) to determine the number of total cells in each sample. Alternatively, OD550 can be used to estimate the number of cells in each sample based on a previously determined standard curve (e.g., count 1 sample and perform serial dilutions to make the curve). Because tumor samples often contain a lot of cellular debris and dead cells which can potentially bind antibodies in subsequent steps, it is important to include them in your total cell estimate.
19. An unstimulated control is used for each sample to determine baseline expression of cytokines/chemokines. Depending on the amount of available sample, questions of interest, and the expression of your analyte, you can opt to not have an unstimulated control for each sample and just rely on the stimulated sample; in this case, you can have a single unstimulated control with a mixture of your samples to determine whether the stimulation worked.
20. It is recommended to have one “cytometer setup “ sample treated the same as other samples to ensure correct settings of the cytometer during acquisition (this prevents the waste of experimental samples during set-up). Fluorescent minus one (FMO) controls are important to have for the RNA targets due to background signal (see Araya & Goldszmid, 2020 for additional information about FMO controls). Have at least one FMO for each RNA analyte using a stimulated sample. If sample groups vastly differ (e.g., different tumors, EL4 vs B16-F10), it is recommended to have separate controls for each group (unstained and FMO).
21. It is crucial to check the plate after each centrifuge step to ensure that there is a clear and intact pellet for each sample. If a clear pellet does not appear, repeat centrifugation step.
22. It is recommended to make small aliquots of the reagent used for stimulation to prevent multiple freeze-thaw cycles. E.g., aliquots of c-di-AMP 2.5 mg/mL in PBS kept at -20°C for use.
23. A U-bottom plate is recommended for the incubation step to increase area for cell contact with c-di-AMP. A V-bottom plate is used for all other steps with centrifugation and staining to ensure cells are pelleted properly.
24. To prevent evaporation of media, sample plate may be placed inside a small receptacle with water in the bottom and a loosely attached lid before placing inside the incubator to increase humidity around the plate.
25. The incubation time can vary and should be adjusted depending on the molecule used for stimulation and the target mRNA. Generally, incubation times can be shorter than those typically used for detection of intracellular cytokine proteins by fluorochrome-conjugated antibodies since the production of mRNA occurs faster. We found that 3 hours was sufficient for the detection of *Ifnb1* and stimulation of the chemokines *Xcl1* and *Ccl5*.
26. When preparing reagent mixes for all samples, it is recommended to add at least 3 extra samples to calculations to account for pipetting error.
27. It is important to perform multiple washes to remove residual FBS that will interfere with the cell viability staining.
28. RNA wash buffer is provided in a large volume bottle in the PrimeFlow™ RNA Assay Kit. It is recommended to make and store aliquots into smaller tubes (e.g., 50 mL) to reduce the amount of time to reach room temperature.
29. The dilution of the antibodies for reference controls may need to be optimized for your sample and cytometer settings. Ideally, reference signal should be within the linear range of your cytometer and at the same or higher intensity than your sample staining. 1 µL of 1:10 diluted antibody stained in a volume of 100 µL (final dilution 1:1000) is given as a suggested starting point.
30. According to manufacturer’s protocol, stock of LD Blue is made by adding 50 µL of provided DMSO to each tube. It is recommended to make small aliquots (<10 µL) and store at -20°C for use to reduce multiple freeze-thaw cycles.
31. The PrimeFlow processing can cause cell pellets to be loose after centrifugation. Manual aspiration using a multichannel pipette is strongly recommended instead of flicking plate upside down to limit cell loss. See Figure 4 for pictures showing loose pellet and the amount of residual liquid desired after supernatant is aspirated.
32. While we have not performed intracellular staining of fluorochrome-conjugated antibodies in conjunction with the PrimeFlow assay, the manufacturer states that it is compatible with the protocol.
33. If desired, the fixation 2 step can be performed by incubating for 60 min at room temperature in the dark before proceeding with the next steps. However, it is recommended to stop here for the day and finish the rest of the staining the following morning.
34. In this protocol, the manufacturer’s protocol for the PrimeFlow RNA Assay Kit in 96-well plates has been adapted in the following ways: reduced amount of key reagents by half (doubled the usage of the kit); reduced number of washes to limit cell loss; and use of a thermocycler with PCR tubes to maintain an incubation temperature of 40±1°C when an accurate incubator with a heat block is not available. All sample manipulation steps are to be performed at room temperature and not on ice.
35. Using the appropriate Target Probes (i.e., Type 1 probe for Xcl1, Type 6 probe for Ccl5, Type 10 probe for Ifnb1), make a Target Probe mix for all samples excluding Unstained and FMOs. For FMOs, make separate mixes containing 2/3 Target Probes for each combination to have an FMO control for each Target Probe.
36. It is important that the residual volume after all washes does not exceed 10 µL. Double check supernatant volumes after pipetting, switch to single pipette if necessary to remove leftover supernatant in specific wells.
37. PCR tubes in conjunction with a thermocycler is used to maintain an accurate temperature of 40±1°C. However, an accurate incubator (40±1°C) with a heat block can be used with a 96-well plate if available.
38. An adapter for PCR tubes may be needed for use in a benchtop centrifuge with swinging buckets. If so, one suggestion is to use two old 200 µL pipette tip racks nested and taped together to make a rack capable of holding the 0.2 mL PCR tubes without falling over.
39. This is an optional safe stopping point according to manufacturer’s instructions; samples can be stored overnight at 4°C in the dark. However, it is recommended to finish RNA staining steps in the same day.
40. Single-color reference controls can be prepared the same day as samples or anytime between then and sample acquisition.
41. 4% PFA (Paraformaldehyde) is made by diluting 16% PFA stock 1:4 in PBS before storing at 4°C in the dark.
42. Cytometer setup will depend on the specific cytometer to be used and the guidelines provided by the manufacturer and/or the cytometer core facility. This may include but is not limited to turning on the machine, refilling sheath fluid, discarding waste, cleaning, and running standardization beads (e.g. CS&T, SpectroFlo QC Beads). It is recommended have pre-set application settings that optimizes channel voltages according to the background of cellular samples (Araya & Goldszmid, 2020).
43. Filter plate can be washed and reused for future experiments. Soak used filter plates in an enzymatic detergent solution (#2252, Advanced Sterilization Products) (see Note 1) for approximately 1 hour followed by thorough rinsing with de-ionized water and air drying overnight. As an alternative to filter plates, samples can be filtered with manually cut 50 µM mesh (#SEFA03-50/31, Sefar) (see Note 1) or 5 mL test tubes with cell-strainer caps (#352235, Falcon) (see Note 1).
44. Choose the most appropriate sample acquisition receptacle for your available cytometer based on previous experience acquiring samples with high amounts of debris. In our experience, cluster tubes set inside 5 mL test tubes worked best for BD instruments such as FACSymphony and LSRFortessa whereas a 96-well plate can be used with instruments using a pressurized fluidics system such as the Cytek Aurora.
45. Tumor samples tend to have a lot of debris/dead cells which can clog fluidics and interfere with acquisition. It is recommended to run <10,000 events/sec and to acquire 1-2 million events to have enough Live CD45^+^ cells for downstream analyses.
46. Use an N x N plot to visualize each data plotted for each pair of markers. Due to the large number of debris and dead cells present in tumor samples, it is recommended to visualize and adjust compensation values on Singlets followed by the Live and Leukocyte gates. Use control samples such as Unstained and FMO as needed to aide in the determination of positive and negative signals.
47. The background signal for each RNA marker will vary and depend on the specific cell type. For example, eosinophils are more autofluorescent leading to a background negative signal that is higher than the positive signal for lymphocytes in some cases. Be careful to define positive and negative signal for each cell type and use the FMO sample.

**Figure 4.**
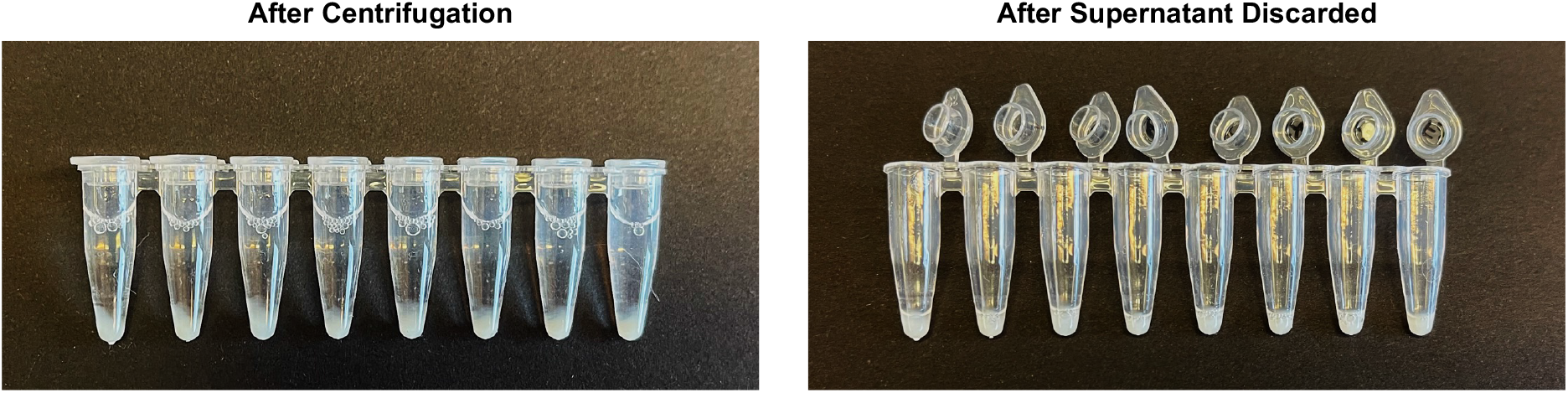
Pictures of cell pellets in PCR tubes during RNA flow cytometry protocol after centrifugation (left) and after supernatant has been discarded manually using a multichannel pipette (right).

## Acknowledgments

We are grateful for the use of the NCI LGI Flow Cytometry Core facility. We thank members of the Goldszmid laboratory for their technical help and feedback. This research was supported by the Intramural Research Program of the NIH (CCR-NCI).

